# Effects of mutating α-tubulin lysine 40 on sensory dendrite development

**DOI:** 10.1101/212373

**Authors:** Brian V. Jenkins, Harriet A. J. Saunders, Helena L. Record, Dena M. Johnson-Schlitz, Jill Wildonger

## Abstract

Microtubules are essential to neuronal structure and function. Axonal and dendritic microtubules are enriched in post-translational modifications that impact microtubule dynamics, transport, and microtubule-associated proteins. Acetylation of α-tubulin lysine 40 (K40) is a prominent, conserved modification of neuronal microtubules. However, the cellular role of microtubule acetylation remains controversial. To resolve how microtubule acetylation might affect neuronal morphogenesis we mutated endogenous α-tubulin in vivo using a new fly strain that facilitates the rapid knock-in of designer *α-tubulin* alleles. Leveraging our new strain, we found that microtubule acetylation, as well as polyglutamylation and (de)tyrosination, is not essential for survival. However, we found that dendrite branch refinement in sensory neurons relies on α-tubulin K40. Mutagenesis of K40 reveals moderate yet significant changes in dendritic lysosome transport, microtubule polymerization, and Futsch distribution in dendrites but not axons. Our studies point to an unappreciated role for α-tubulin K40 and acetylation in dendrite morphogenesis. While our results are consistent with the idea that microtubule acetylation patterns microtubule function within neurons, they also suggest there may be a structural requirement for α-tubulin K40.

**Summary Statement:** Neurons are enriched in post-translationally modified microtubules. Targeted mutagenesis of endogenous α-tubulin in flies reveals that dendrite branch refinement is altered by acetylation-blocking mutations.

## INTRODUCTION

Microtubules provide the basis for neuronal architecture. The ability of neurons to transmit and receive signals depends on the proper morphogenesis of axons and dendrites. Axons and dendrites differ in structure as well as function. Microtubules in each compartment are uniquely organized and enriched in post-translational modifications (PTMs), including acetylation, detyrosination, and polyglutamylation (Chakraborti et al., 2016; Song and Brady, 2015). The patterns of microtubule PTMs between and within axons and dendrites are thought to be critical to functional compartmentalization by locally regulating microtubule dynamics and/or transport. Yet the role that microtubule PTMs, in particular acetylation, may play in neuronal morphogenesis has been controversial.

Several conserved lysines in α- and β-tubulin are acetylated, and acetylation of the α-tubulin luminal residue lysine 40 (K40) has been the most well-studied since its discovery over thirty years ago (Choudhary et al., 2009; Chu et al., 2011; Howes et al., 2014; L'Hernault and Rosenbaum, 1983; L'Hernault and Rosenbaum, 1985; Soppina et al., 2012). Acetylation of α-tubulin K40 was initially characterized as a marker of microtubules resistant to depolymerizing drugs (Piperno et al., 1987). Although acetylation typically correlates with stable, long-lived microtubules in cells, acetylation itself does not confer stability, but rather may make microtubules more resilient to mechanical forces as microtubules age (Coombes et al., 2016; Howes et al., 2014; Ly et al., 2016; Palazzo et al., 2003; Portran et al., 2017; Szyk et al., 2014; Webster and Borisy, 1989; Wilson and Forer, 1997; Xu et al., 2017). Yet despite years of study, the effects of acetylation on microtubules and microtubule function in cells are still debated.

In cultured mammalian neurons, young axons are enriched in acetylated microtubules in comparison to dendrites. This difference initially led to the idea that acetylation might label microtubule tracks for selective transport to one compartment or the other (Song and Brady, 2015). Consistent with this idea, acetylation has been shown to distinguish the microtubule tracks that are preferentially bound by kinesin-1, which transports cargo from the cell body to axon terminal (Dompierre et al., 2007; Guardia et al., 2016). The neuron-wide expression of α-tubulin K40Q, which mimics acetylation, has also been reported to redirect kinesin-1 to dendrites (Farias et al., 2015). Similarly, in immature unpolarized neurons, increasing microtubule acetylation redirects kinesin-1 to multiple neurites (Hammond et al., 2010; Reed et al., 2006). However, in mature polarized neurons, microtubule acetylation by itself is not sufficient to alter kinesin-1 localization (Cai et al., 2009; Hammond et al., 2010). Also, microtubule acetylation does not affect kinesin-1 motility in purified in vitro systems (Kaul et al., 2014; Walter et al., 2012). Thus, there are conflicting reports about whether microtubule acetylation is necessary and/or sufficient to affect motor activity and localization in neurons.

There is also conflicting evidence regarding the role of microtubule acetylation in neuronal development. The function of microtubule acetylation in the developing nervous system has been investigated mainly through the loss or over-expression of the primary α-tubulin acetyltransferase and deacetylase enzymes, αTAT1 and HDAC6, respectively (Akella et al., 2010; Hubbert et al., 2002; Shida et al., 2010). On one hand, there are reports that inhibiting HDAC6 disrupts axon initial segment formation in cultured neurons (Tapia et al., 2010; Tsushima et al., 2015), and that cortical neuron migration is impeded by either the knock-down of αTAT1 or the over-expression of α-tubulin K40A, which cannot be acetylated (Creppe et al., 2009; Li et al., 2012). On the other hand, *HDAC6* and *αTAT1* knock-out mice are homozygous viable. Neither knock-out results in any gross neurological defect, such as a disruption in cortical layering, that is typically associated with abnormal neuronal polarity (Kalebic et al., 2013; Kim et al., 2013; Zhang et al., 2008). Worms lacking αTAT1 activity are viable, but touch insensitive (Akella et al., 2010; Cueva et al., 2012; Shida et al., 2010; Topalidou et al., 2012; Zhang et al., 2002). A recent study has shown that *αTAT1* knock-out mice are also insensitive to mechanical touch and pain (Morley et al., 2016), indicating that the functional effects of microtubule acetylation are likely conserved between invertebrates and vertebrates. These functional studies raise the question of whether and how microtubule acetylation might sculpt neuronal architecture. Here again, there is conflicting evidence arguing both for and against the importance of acetylated microtubules to axonal morphology (Morley et al., 2016; Neumann and Hilliard, 2014). It is not known whether only axons rely on acetylated microtubules; indeed, a potential role for microtubule acetylation in dendrite morphogenesis has not been explored.

We sought to resolve the role of microtubule acetylation in neuronal transport and morphogenesis through targeted mutagenesis of endogenous α-tubulin in Drosophila. A key advantage of mutating endogenous α-tubulin is that we can directly and specifically assess the involvement of α-tubulin residues in the development of axons and dendrites as well as microtubule growth and microtubule-dependent activities. Our approach leverages a new fruit fly strain that we created to enable the rapid knock-in of designer *α-tubulin* alleles. By directly targeting the α-tubulin residues that are modified, we avoid complications often associated with targeting the modifying enzymes. For example, several modifying enzymes have cellular targets in addition to tubulin. While this is not the case for αTAT1, which acetylates itself and α-tubulin K40, HDAC6 deacetylates multiple proteins in addition to α-tubulin (Valenzuela-Fernandez et al., 2008). Some enzymes, such as polyglutamylases, can modify several tubulin residues and some modifying enzymes, such as the carboxypeptidase that removes the terminal tyrosine on α-tubulin, remain unidentified (Janke, 2014; Song and Brady, 2015). This presents challenges to using an enzyme-based approach to dissect the role of microtubule PTMs in cells.

Through live imaging of sensory neurons in developing fruit flies, we found that targeted mutagenesis of endogenous α-tubulin K40 does not disrupt selective transport to axons or dendrites, or neuronal polarity, but does affect the refinement of dendrite branches. Acetylation-blocking mutations increase branch number with a correlative increase in terminal branch growth. Both α-tubulin K40A and K40R mutations block acetylation. However, only the arginine substitution conserves the length and charge of the lysine sidechain; alanine does not and thus may alter α-tubulin structure. We found that the K40R mutation does not phenocopy the effects of the K40A mutation on dendrite dynamics, suggesting that K40 may be essential for α-tubulin and/or microtubule structure. In the α-tubulin K40A mutant dendrites we observed modest yet significant changes in lysosome transport, microtubule growth, and Futsch distribution that might underlie an increase in branch number. Combined, our data point to a previously unappreciated role for K40 and acetylation in fine-tuning dendrite patterning.

## RESULTS

### Characterization of α-tubulin mutations that disrupt microtubule PTMs

To determine the function of microtubule PTMs in neurons *in vivo*, we undertook targeted mutagenesis of α-tubulin in fruit flies. Like other organisms, the *Drosophila melanogaster* genome has several distinct α-tubulin genes that encode unique protein isotypes, which assemble into microtubules that are modified. The four Drosophila α-tubulin genes have been named based on their cytological location: *αTub84B*, *αTub84D*, *αTub85E*, and *αTub67C* (Raff, 1984). αTub84B is likely the predominant α-tubulin in flies and is 97% *identical* to human TUBA1A, with only five non-conservative amino acid differences, four of which are within the C-terminal tail (Fig. 1A). Like α-tubulin in other organisms, αTub84B is modified at multiple residues (Bobinnec et al., 1999; Piperno and Fuller, 1985; Warn et al., 1990; Wolf et al., 1988). In the sensory class IV dendritic arborization (da) neurons that we use as a model, microtubules are acetylated, polyglutamylated, and tyrosinated (Fig. 1B-F and data not shown). In embryos, axonal microtubules are heavily acetylated (Fig. 1C,D), consistent with findings that young axons of mammalian neurons in culture are also enriched in acetylated microtubules. In mature larval da neurons, microtubule acetylation levels are equivalent between axons and dendrites (Fig. 1E-F).

**Fig. 1.**
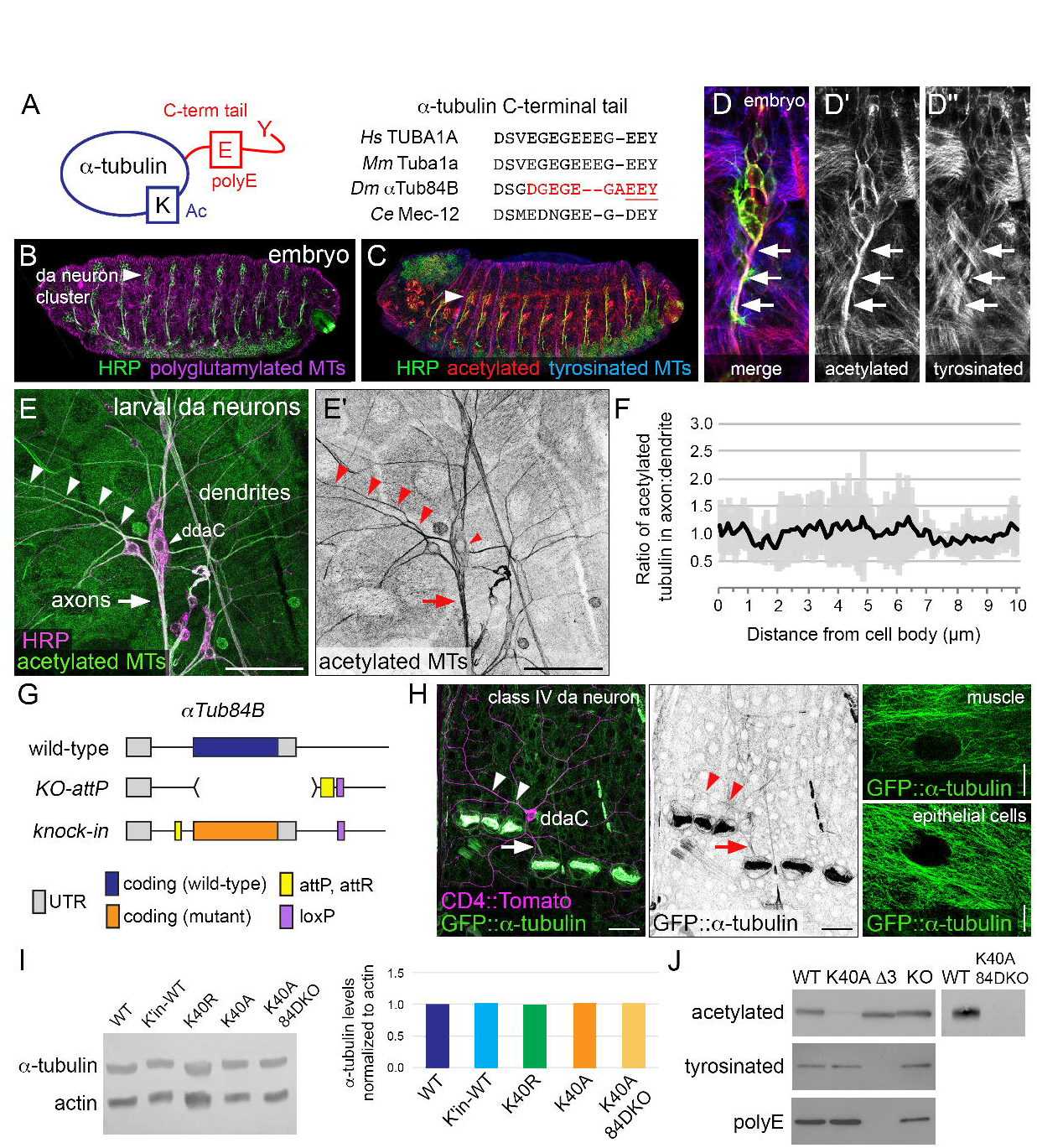
*In vivo* analysis of α-tubulin modifications. (A) Cartoon showing α-tubulin modifications (left) and sequence alignment of the C-terminal tails of human, mouse, fly, and worm Tuba1a orthologs (right). Red: ten amino acids in αTub84B^ΔC^. Underline: three residues (EEY) deleted in αTub84B^Δ3^. (B-D) Microtubules in developing fly embryos are modified and da neuron axons are enriched in acetylated microtubules (D-D"). Embryos were stained for HRP (green), a neuronal membrane marker, as well as polyglutamylated (magenta, B) acetylated (red, C and D), and tyrosinated (blue, C and D) microtubules. Scale bars: 100 µm (B) and 10 µm (D). Arrowheads: da neuron cluster; arrows: axons. (E-F) In larval da neurons, microtubules in axons and dendrites are acetylated at equivalent levels. The acetylated microtubule signal in proximal axonal and dendritic segments were compared as a ratio (mean ± SD), n = 7 class IV ddaC neurons. Green: acetylated microtubules and magenta: HRP. Scale bar: 50 µm. Arrowheads: dendrite and ddaC cell body marker; arrow: axons. Images from fixed tissue. (G) Cartoon of the *αTub84B*^*KO-attP*^ allele that enables rapid knock-in of designer alleles to interrogate α-tubulin function *in vivo*. Top: wild-type *αTub84B* (blue), middle: the major coding exon of *αTub84B* was deleted (brackets) and replaced by an *attP* site (yellow), bottom: knock-in of a mutant *αTub84B* (orange). (H) GFP-tagged αTub84B is broadly expressed in developing larvae. In muscles and epithelial cells, a filamentous pattern indicates GFP::αTub84B is likely incorporated into microtubules. Scale bars: 25 µm (left) and 10 µm (muscle and epithelial cell images, right). Arrowheads: dendrite; arrow: axon. Green: GFP, magenta: CD4::Tomato. Images from live 3^rd^ instar larvae. (I) Western blot analysis (left) and quantification (right) of α-tubulin levels (normalized to actin) in mutant strains as indicated. All strains are homozygous. K40A 84DKO refers to *αTub84B*^*K40A*^ chromosome with *αTub84D* deleted *(αTub84B^K40A^, αTub84D*^*KO*^). (J) Western blot of lysates from wild-type and *αTub84B* mutant fly heads probed for acetylated, tyrosinated, and polyglutamylated α-tubulin, as indicated. The two-lane blot (right) was probed for acetylated α-tubulin includes lysate from double-mutant *αTub84B*^*K40A*^ *αTub84D*^*KO*^ fly heads. The αTub84B Δ3 mutation eliminates both the anti-tyrosinated and anti-polyglutamylated α-tubulin signals.

Using a genome-endendrites of control andgineering approach, we created a new fly strain that enables us to readily knock-in designer *αTub84B* alleles via site-directed recombination (Fig. 1G). We used an ends-out gene targeting strategy (Huang et al., 2009) to replace *αTub84B* with an *attP* "landing" site. Consistent with previous reports, deleting *αTub84B* resulted in lethality (Table 1), indicating that *αTub84B* is essential for survival and that the other α-tubulin genes could not compensate for its loss (Matthews and Kaufman, 1987). This includes *αTub84D*, which has a similar expression pattern and encodes a nearly identical protein that differs from αTub84B by two amino acids (Matthews et al., 1989; Raff, 1984). The knock-out strain was rescued by knocking-in wild-type *αTub84B* (*αTub84B*^*K'in-WT*^), indicating that the *attP* replacement strategy did not disrupt the function of the *αTub84B* locus (Table 1). To confirm that *αTub84B* is indeed broadly expressed, including in the nervous system (Raff, 1984), we generated flies that express GFP-tagged αTub84B. As expected, GFP:: αTub84B was expressed in most cell types, including neurons (Fig. 1H). In muscles and epithelial cells, GFP:: αTub84B appeared filamentous, suggesting GFP-tagged tubulin was incorporated into microtubules (Fig. 1H). However, it should be noted that the *GFP::aTub84B* allele is dominant male sterile and does not survive in trans to a deletion that removes *aTub84B*. This suggests that GFP::aTub84B does not function equivalently to the wild-type untagged protein. Thus, we have created a unique and powerful tool to manipulate and visualize endogenous α-tubulin *in vivo*.

**Table 1.** Effects of α-tubulin mutations on survival.

| *α-tubulin* alleles | Description of mutation | Effect on survival |
| --- | --- | --- |
| *αTub84B*<sup>KO</sup> | deletes *αTub84B* | lethal |
| *αTub84B*<sup>K'in-WT</sup> | knock-in wild-type *αTub84B* | viable |
| *GFP::αTub84B* | knock-in GFP-tagged wild-type *αTub84B* | lethal* |
| *αTub84B*<sup>ΔC</sup> | last 10 aa of C-terminal tail (CTT) deleted | lethal* |
| *αTub84B*<sup>Tuba1a-tail</sup> | fly CTT replaced by vertebrate CTT | viable |
| *αTub84B*<sup>Δ3</sup> | last 3 aa of CTT deleted, blocks (de)tyrosination | viable |
| *αTub84B*<sup>AAAA</sup> | 4 glutamates in CTT mutated, blocks polyglutamylation and glycylation of CTT | viable |
| *αTub84B*<sup>K40A</sup> | blocks K40 acetylation (charge and length of lysine sidechain not conserved) | viable |
| *αTub84B*<sup>K40R</sup> | blocks K40 acetylation (charge and length of lysine sidechain conserved) | viable |
| *αTub84B*<sup>K40A</sup>,<br>*αTub84D*<sup>KO</sup> | blocks *αTub84B* K40 acetylation and deletes *αTub84D* | viable<sup>‡</sup> |

\*dominant male sterile and lethal in trans to deletion strains that eliminate *$\alpha$ Tub84B*.
<sup>‡</sup> *$\alpha$ Tub84B*<sup>K40A</sup>, *$\alpha$ Tub84D*<sup>KO</sup> is viable in trans to deficiencies that eliminate *$\alpha$ Tub84D*
(Df(3R)BSC747, Df(3R)ED7665, Df(3R)BSC465) and to a deficiency that removes both
*$\alpha$ Tub84B* and *$\alpha$ Tub84D* (Df(3R)Antp17).

We targeted K40 acetylation as well as two additional α-tubulin modifications that have also been implicated in neuronal development and transport, namely polyglutamylation and detyrosination of the C-terminal tail. Many of the studies on microtubule polyglutamylation and (de)tyrosination have been carried out in vertebrate models. The fly and mammalian C-terminal tails differ in several amino acids, including several glutamate residues that are sites of polyglutamylation (Fig. 1A). We tested whether the function of the α-tubulin C-terminal tails from these different species might be conserved despite the sequence differences. Replacing the αTub84B C-terminal tail with that of the mammalian Tuba1a did not affect viability (Table 1), indicating the mammalian Tuba1a C-terminal tail can functionally substitute for the fly αTub84B C-terminal tail (Fig. 1A). Deletion of the C-terminal tail resulted in lethality (Table 1), indicating that the C-terminal tail is essential for proper α-tubulin function in vivo. Blocking two different modifications of the C-terminal tail, polyglutamylation *(αTub84B*^*AAAA*^) and detyrosination-tyrosination *(αTub84B*^*Δ3*^), did not affect animal survival (Table 1). Since the glutamates in the C-terminal tail are thought to mediate interactions with essential motors and other microtubule-binding proteins, it was particularly surprising that eliminating virtually all the negatively charged residues in *αTub84B*^*AAAA*^ did not affect viability (Bonnet et al., 2001; Boucher et al., 1994; Lacroix et al., 2010; Larcher et al., 1996; Roll-Mecak, 2015; Sirajuddin et al., 2014; Valenstein and Roll-Mecak, 2016; Wang and Sheetz, 2000). Combined, our results suggest that the C-terminal tail has a conserved role in α-tubulin function *in vivo*, yet polyglutamylation and modification of the terminal residues of the C-terminal tail are dispensable for survival.

To test the role of αTub84B K40 acetylation in survival and neuronal morphogenesis we introduced K40A and K40R mutations to eliminate acetylation. Both *αTub84B* alleles were viable and fertile in trans to the *αTub84B* knock-out (Table 1), consistent with reports that loss of K40 acetylation does not affect survival (Akella et al., 2010; Cueva et al., 2012; Kalebic et al., 2013; Kim et al., 2013; Mao et al., 2017; Shida et al., 2010; Topalidou et al., 2012; Zhang et al., 2008). Our viability and fertility results agree with a recent study that used CRISPR-Cas9 to introduce the K40R mutation into *αTub84B* (Mao et al., 2017). Western blot analysis of adult fly head lysate revealed that the amounts of the mutant αTub84B proteins were equivalent to wild-type and that α-tubulin K40 acetylation was virtually abolished in the *αTub84B*^*K40A*^ flies (Fig. 1I,J). The residual signal in the western blot may reflect acetylation of another α-tubulin isotype, most likely αTub84D, which is also broadly expressed (Raff, 1984). We used CRISPR-Cas9 genome editing to delete the entire αTub84D gene in the *αTub84B*^*K40A*^ strain. The *αTub84B*^*K40A*^ *αTub84D*^*KO*^ double mutant eliminated the residual acetylated tubulin signal in western blots (Fig. 1J) and was viable in trans to a large deletion that removes both α-tubulin genes (Table 1). Genetic complementation tests also unexpectedly revealed that *αTub84D* is a non-essential gene (Table 1). Combined, these data indicate α-tubulin K40 acetylation is not essential for survival.

### αTub84B K40A does not affect selective transport to axons, but has a compartment-specific effect on retrograde lysosome transport in dendrites

Microtubule acetylation has been shown to affect microtubule-based transport in cultured cells, including neurons (Dompierre et al., 2007; Reed et al., 2006). One recent model suggests that acetylated microtubules are part of an exclusion zone that prevents dendritic cargos from entering axons (Farias et al., 2015). Yet there is also evidence that microtubule acetylation alone is not sufficient to direct motors to a specific compartment (Atherton et al., 2013). The class IV da neurons that we use as a model reside just below the transparent larval cuticle, allowing for live imaging of transport in neurons in intact animals. First, we examined the distribution of a polarized organelle population, Golgi outposts, which localize to dendrites and regulate dendrite patterning in flies and mammals (Horton and Ehlers, 2003; Horton et al., 2005; Ye et al., 2007). We found that the polarized dendritic localization of Golgi outposts was not altered in *αTub84B*^*K40A*^ neurons (Fig. 2A-B'''). Thus, microtubule acetylation is not an essential part of the mechanism that prevents Golgi outposts from entering axons.

**Fig. 2.**
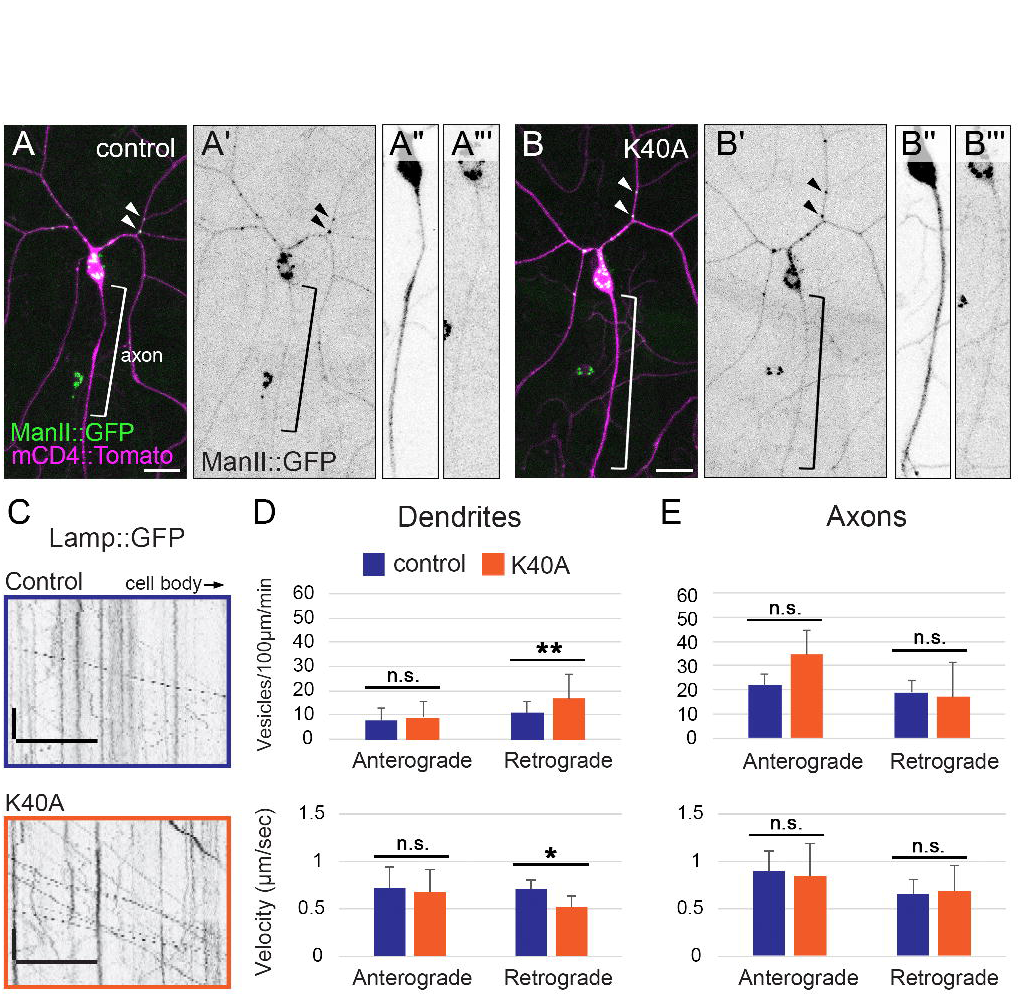
The αTub84B K40A mutation does not affect the polarized distribution of Golgi outposts, but does affect lysosome motility in dendrites. (A-B''') Golgi outposts, marked by ManII::GFP (green), localize to dendrites in both control (A-A''') and *αTub84B*^*K40A*^ neurons (B-B'''). ManII::GFP (green in A and B, black in A', A''', B', and B''') and CD4::Tomato (magenta in A and B, black in A'' and B'') are expressed in class IV da neurons under the control of the *ppk* enhancer. Bracket: axon, arrowheads: Golgi outposts in dendrites. Scale bar: 25 µm. (C) Representative kymographs of lysosome dynamics in the dendrites of control (top) and *αTub84B*^*K40A*^ (bottom) neurons. Scale bar x-axis: 10 µm, scale bar y-axis: 10 sec. Lysosomes are marked by Lamp1::GFP. Cell body is to the right. (D,E) In dendrites (D), lysosomes traveling retrograde in *αTub84B*^*K40A*^ neurons display increased flux (top) and reduced velocity (bottom). Lysosome motility in axons (E) is unaffected by the αTub84B K40A mutation. Dendrites (D, flux): 30 wild-type control dendrite segments and 29 *αTub84B*^*K40A*^ dendrite segments were analyzed (mean ± SD); p=0.008. Dendrites (D, velocity): 32 wild-type control dendrite segments and 29 *αTub84B*^*K40A*^ dendrite segments were analyzed (mean ± SD); p=0.012. Axons (E, flux): 7 wild-type control axons and 5 *αTub84B*^*K40A*^ axons were analyzed (mean ± SD). Axons (E, velocity): 7 wild-type control axons and 5 *αTub84B*^*K40A*^ axons were analyzed (mean ± SD). Statistical significance was evaluated using a two-tailed Student’s t-test. **p=0.001-0.01, *p=0.01-0.05, n.s. = not significant.

Although selective transport to dendrites or axons is not perturbed, it is possible that microtubule acetylation affects other aspects of microtubule-based transport. The trafficking of lysosomes, an organelle component of the autophagy pathway, is sensitive to microtubule acetylation in cultured cells (Chauhan et al., 2015; Guardia et al., 2016; Xie et al., 2010). Notably, a recent study revealed that microtubule acetylation distinguishes a set of tracks that are preferentially used by kinesin-1 to transport lysosomes in the perinuclear region in HeLa cells (Guardia et al., 2016). We expressed the lysosome marker Lamp1::GFP in the da neurons and analyzed its dynamic localization in the axons and dendrites of control and *αTub84B*^*K40A*^ neurons. In *αTub84B*^*K40A*^ dendrites, the velocity of lysosomes traveling retrogradely was significantly reduced while their flux nearly doubled (Fig. 2C,D). Lysosomes moving anterogradely in *αTub84B*^*K40A*^ dendrites were not affected (Fig. 2D). Since microtubules in da neuron dendrites are oriented predominantly minus-end-distal (Rolls et al., 2007), lysosomes moving anterogradely are likely transported by dynein whereas those that move retrogradely are likely transported by kinesin. In contrast, lysosome transport in axons was unchanged in the *αTub84B*^*K40A*^ neurons (Fig. 2E). Our data indicate that the αTub84B K40A mutation selectively alters the retrograde, likely kinesin-mediated, transport of lysosomes in dendrites, but does not affect either the retrograde or anterograde transport of lysosomes in axons. While the αTub84B K40A mutation does not affect selective transport to axons or dendrites, it does have a dendrite-specific effect on lysosome transport.

### αTub84B K40A and K40R mutations increase dendrite branch number

The axons of young neurons in culture are enriched in acetylated microtubules relative to dendrites (Song and Brady, 2015), which has led to a focus on the role of microtubule acetylation in axons. However, our analyses of lysosome transport in *αTub84B*^*K40A*^ neurons suggest that dendritic, not axonal, microtubules are sensitive to α-tubulin K40 mutagenesis. We next tested whether dendrite morphogenesis is affected by K40 mutations. In developing larvae, the class IV da neurons extend an expansive dendritic arbor over a shallow depth, making them ideal for analyzing dendrite growth and patterning (Grueber et al., 2002). Acetylated microtubules are present in the main dendrite branches and a subset of terminal branches of class IV da neurons (Fig. 3A). To visualize dendrite arbors and quantify dendrite branching in control and αTub84B K40 mutants, we used the transgene *ppk-CD4::GFP*, which expresses a GFP-tagged transmembrane protein (CD4) under the control of the class IV-specific *pickpocket* (*ppk*) enhancer. The overall dendritic coverage of the *αTub84B*^*K40A*^ and *αTub84B*^*K40R*^ neurons appears normal in that the mutant arbors extended to the segment boundaries normally and tiled properly with their neighbors. However, our quantification of dendrite tips revealed that both the K40A and K40R mutations resulted in an increased number of terminal branches compared to age-matched wild-type controls at 120 hours (h) after egg laying (AEL) (Fig. 3B-E). We did not detect any defects in axon termination in the ventral nerve cord (data not shown).

**Fig. 3.**
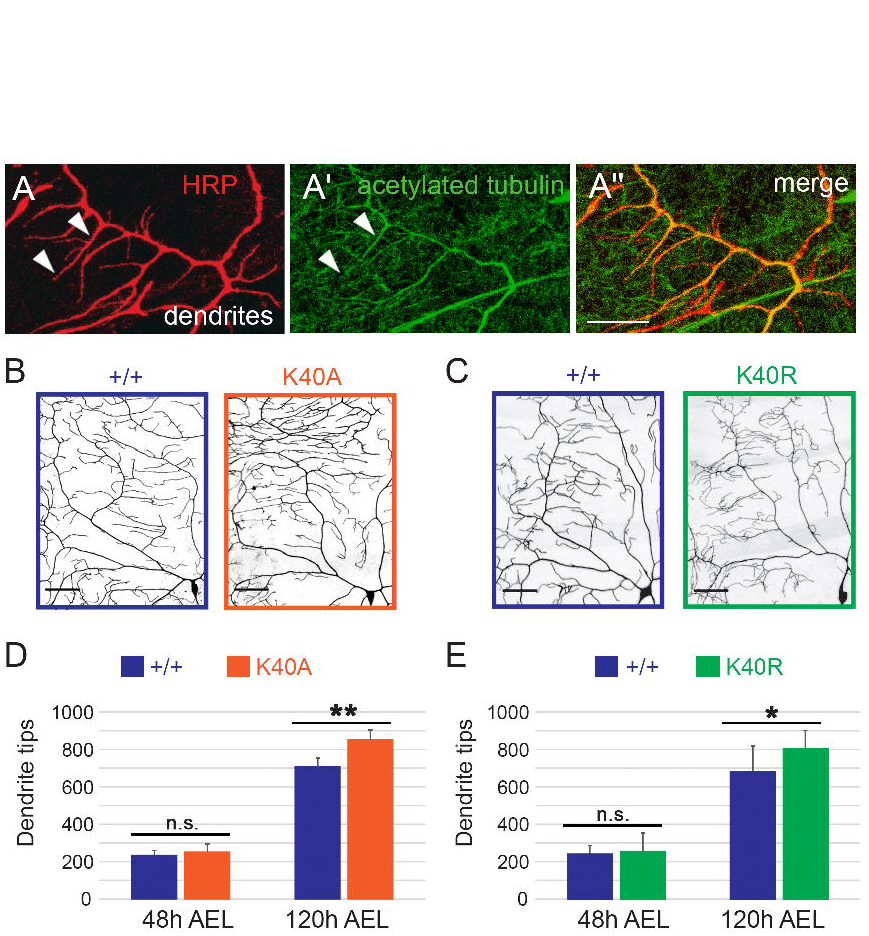
Sensory dendrite tip number is increased in αTub84B K40 mutant neurons. (A) In class IV da neuron dendrites, acetylated microtubules are present in main dendrite branches and some terminal dendrites (arrowheads). Red: HRP, neuronal membrane marker, green: anti-acetylated α-tubulin. Scale bar: 10 µm. (B-E) Mutations that prevent αTub84B K40 acetylation (K40A, K40R) increase the number of dendrite tips at 120 h AEL. Terminal branch number is not significantly affected at 48 h AEL. Quadrant of class IV da neuron dendrite arbor at 120 h AEL illuminated with *ppk-CD4::GFP* (B,C). Scale bar: 50 µm. Experiments to analyze the morphogenesis of each mutant included age-matched controls that were imaged and analyzed in parallel. The numbers of dendrite tips in control neurons do not significantly differ between each other. *αTub84B*^*K40A*^ dendrite analysis (D): 6 wild-type control and 6 *αTub84B*^*K40A*^ neurons were analyzed (mean ± SD), p=0.001. *αTub84B*^*K40R*^ dendrite analysis (E): 8 wild-type control and 14 *αTub84B*^*K40R*^ neurons were analyzed (mean ± SD), p=0.04. Statistical significance was evaluated using one-way ANOVA with post hoc two-tailed Student’s t-tests between experimentally matched control and mutant neurons. **p=0.001-0.01, *p=0.01-0.05, n.s. = not significant.

### Dendrite branch growth is increased in αTub84B K40A mutant neurons

The class IV da neuron dendrites initially extend during late embryonic stages and continue to grow throughout larval stages. Dendrite branches undergo remodeling and refinement through bouts of de novo growth, extension and retraction as larvae grow in size (Parrish et al., 2009; Stewart et al., 2012). During early larval stages (48 h AEL at the beginning of the 2^nd^ larval instar), terminal branches are dynamic and new branches are added to the arbor whereas during late larval stages, terminal branches are less dynamic and fewer new branches appear. At 48 h AEL both K40A and K40R mutant neurons had the same number of dendrite tips as control neurons (Fig. 3B-E), indicating that the increase in dendrite tip number was likely not due to a defect in initial stages of dendrite extension during embryonic stages. Rather, we reasoned that the increase in terminal dendrites might have resulted from changes in dendrite dynamics during early larval stages. To test whether dendrite tip number increased due to increased branch growth and/or decreased branch retraction, we used time-lapse imaging to record dendrite dynamics in 48 h AEL larvae. We then quantified the number of dendrite tips that formed de novo, extended or retracted over a 15-minute interval (Fig. 4A). In the *αTub84B*^*K40A*^ neurons, significantly more branches extended compared to controls, albeit de novo branch growth did not significantly increase over this time interval (Fig. 4 B-D). While *αTub84B*^*K40R*^ mutant neurons had an increased number of dendrite tips like *αTub84B*^*K40A*^, neither dendrite growth nor retraction was significantly altered (Fig. 4B-D). This suggests the increase in dendrite tips in the *αTub84B*^*K40A*^ arbors is likely due to an increase in dendrite growth. Moreover, the difference between dendrite dynamics in the αTub84B K40A versus K40R mutant neurons suggests that K40 may be structurally important for α-tubulin and/or microtubule function in neurons.

**Fig. 4.**
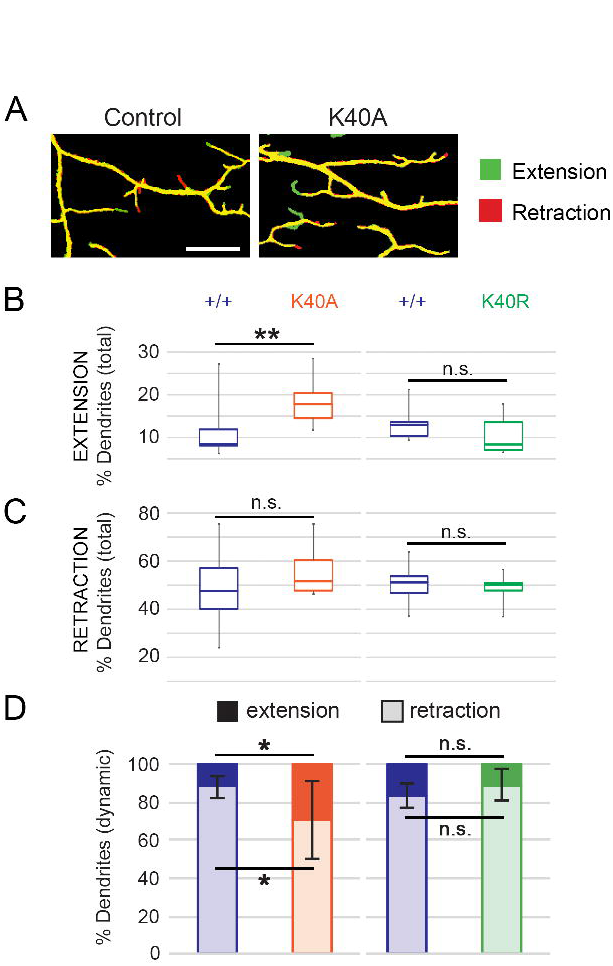
Terminal-dendrite extension is increased in αTub84B^K40A^ neurons. (A) Overlaid photographs of a dendrite imaged at two time points 15 minutes apart in control (left) and *αTub84B*^*K40A*^ (right) larvae at 48 h AEL. The initial dendrite image (t=0) is pseudo-colored red and the second image (t=15 min) is shown in green. Dendrites that have retracted will appear red and those that have extended will appear green. Scale bar: 10 µm. (B,C) Percentage of the total number of terminal branches that extended (B) or retracted (C) during the 15 min imaging interval in wild-type control, *αTub84B*^*K40A*^, and *αTub84B*^*K40R*^ neurons. Boxes represent first and third quartiles (median indicated by line) and whiskers indicate minimum and maximum values. (D) Percentage of the dynamic terminal branches (mean ± SD) that either extended (dark bar) or retracted (light bar). Experiments included age-matched controls that were imaged and analyzed in parallel; all dendrites in the entire arbor of each neuron were analyzed. Dendrite dynamics between control neurons are not significantly different. *αTub84B*^*K40A*^ dendrite analysis (B-D): 12 wild-type control and 8 *αTub84B*^*K40A*^ neurons were analyzed, p=0.002 (% total dendrites that extended), p=0.014 (% dynamic dendrites that extended or retracted). *αTub84B*^*K40R*^ dendrite analysis (B-D): 11 wild-type control and 8 *αTub84B*^*K40R*^ neurons were analyzed. Statistical significance was evaluated using one-way ANOVA with post hoc two-tailed Student’s t-tests between experimentally matched control and mutant neurons. ***p=0.0001-0.001, **p=0.001-0.01, *p=0.01-0.05, n.s. = not significant.

### Dendritic microtubule polymerization frequency is reduced by αTub84B K40A

Next, we analyzed whether the increase in dendrite growth in the αTub84B K40A mutants might reflect a change in the growth of the microtubules themselves. We focused on the αTub84B K40A mutant as it resulted in significantly altered dynamic dendrite growth. Although acetylation does not affect microtubule polymerization in vitro (Dompierre et al., 2007; Howes et al., 2014; Maruta et al., 1986), we nonetheless tested this possibility since microtubule growth has been previously correlated with terminal branch growth in da neurons (Ori-McKenney et al., 2012; Sears and Broihier, 2016; Yalgin et al., 2015). Based on these reports, we predicted that the increase in terminal branch growth in the αTub84B K40 mutants might correlate with an increase in microtubule growth. To monitor microtubule growth we used EB1::GFP, which associates with the plus-ends of growing microtubules in neurites (Fig. 5A). Our data reveal that the αTub84B K40A mutation resulted in a reduced number of EB1::GFP comets specifically in dendrites (Fig. 5B,C) demonstrating that blocking K40 acetylation affected the polymerization of dendritic, but not axonal, microtubules. The rate at which microtubules polymerized was not affected by the K40A mutation (control dendrites: 0.123 ± 0.028 µm min^−1^, n=25, and *αTub84B*^*K40A*^ dendrites: 0.113 ± 0.029 µm min^−1^, n=30, p = 0.19). Thus, similar to its effect on lysosomes, the αTub84B K40A mutation had a compartment-specific effect on microtubule polymerization.

**Fig. 5.**
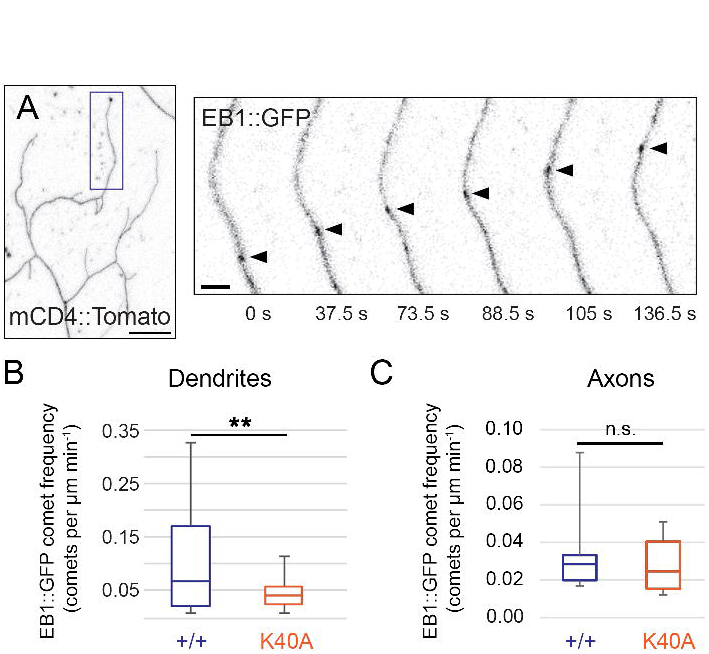
Reduced microtubule growth frequency in dendrites of αTub84B^K40A^ neurons. (A) In a terminal dendrite (box, left panel), EB1::GFP (arrowhead, right panels) marks a microtubule growing towards the dendrite tip in a *αTub84B*^*K40A*^ neuron. Scale bars: 10 µm (left panel) and 2 µm (right panels). (B,C) Microtubule polymerization frequency, quantified as the number of EB1::GFP comets per µm per minute, is significantly decreased in dendrites (B), not axons (C), of *αTub84B*^*K40A*^ neurons. Boxes represent first and third quartiles (median indicated by line) and whiskers indicate minimum and maximum values. Experiments included age-matched controls that were imaged and analyzed in parallel. EB1::GFP analysis: 27 wild-type control and 30 *αTub84B*^*K40A*^ dendrites were analyzed (B), p=0.003; 6 wild-type control and 6 *αTub84B*^*K40A*^ axons were analyzed (C). Statistical significance was evaluated using a two-tailed Student’s t-test. **p=0.001-0.01, *p=0.01-0.05, n.s. = not significant.

### Proximal-distal gradient of Futsch in dendrites is disrupted by αTub84B K40A

We next considered whether mutating K40 might affect dendrite branch growth via an effect on microtubule associated proteins (MAPs), which are also known to regulate dendrite branch growth (Conde and Caceres, 2009). Microtubule acetylation might impact the activity and/or distribution of a MAP that is important for proper dendrite branching. We initially tested whether loss of microtubule acetylation disrupts microtubule severing by katanin. Katanin has been previously shown to be sensitive to microtubule acetylation levels in dendrites (Sudo and Baas, 2010). However, we found that the loss of αTub84B K40 acetylation neither blocks nor enhances katanin-induced changes in dendrite morphogenesis (Fig. S1). This is consistent with a recent report that modulating HDAC6 levels does not affect katanin-induced dendrite growth defects (Mao et al., 2014).

We next turned to Futsch, the fly homolog of MAP1B, which has been shown to regulate dendrite branching (Sears and Broihier, 2016; Yalgin et al., 2015) and has a similar distribution pattern as acetylated microtubules in dendrites (Fig. 6A, Fig. 1E,E' and (Grueber et al., 2002; Jinushi-Nakao et al., 2007). We asked whether the distribution of Futsch might be affected by the acetylation-blocking K40A mutation. Consistent with previous reports (Grueber et al., 2002; Jinushi-Nakao et al., 2007), in wild-type control neurons we found that Futsch was enriched in the main dendrite branches and decreased towards the dendrite tips, where it was detected in some, but not all, terminal branches (Fig. 6A). In control dendrites, Futsch levels decayed ~ 70% from the cell body to distal dendrite tip (Fig. 6B). In *αTub84B*^*K40A*^ neurons, the proximal-medial dendrite segments showed a significant decrease in Futsch, but Futsch levels were comparable to control dendrites in the medial-distal segments (Fig. 6B). Futsch levels in control and *αTub84B*^*K40A*^ axons were equivalent (Fig. 6B). While decreased Futsch has been shown to increase dendrite branch number in one study (Yalgin et al., 2015), another study has found the opposite (Sears and Broihier, 2016). In agreement with the first study, we found that the reduction of Futsch in a *futsch* hypomorph (*futsch*^*K68*^) increased dendrite branch number (Fig. 6C-E). Combined, our results suggest a model in which a change in Futsch distribution in the dendrite arbor may contribute to the increase in dendrite tips in the *αTub84B*^*K40A*^ neurons.

**Fig. 6.**
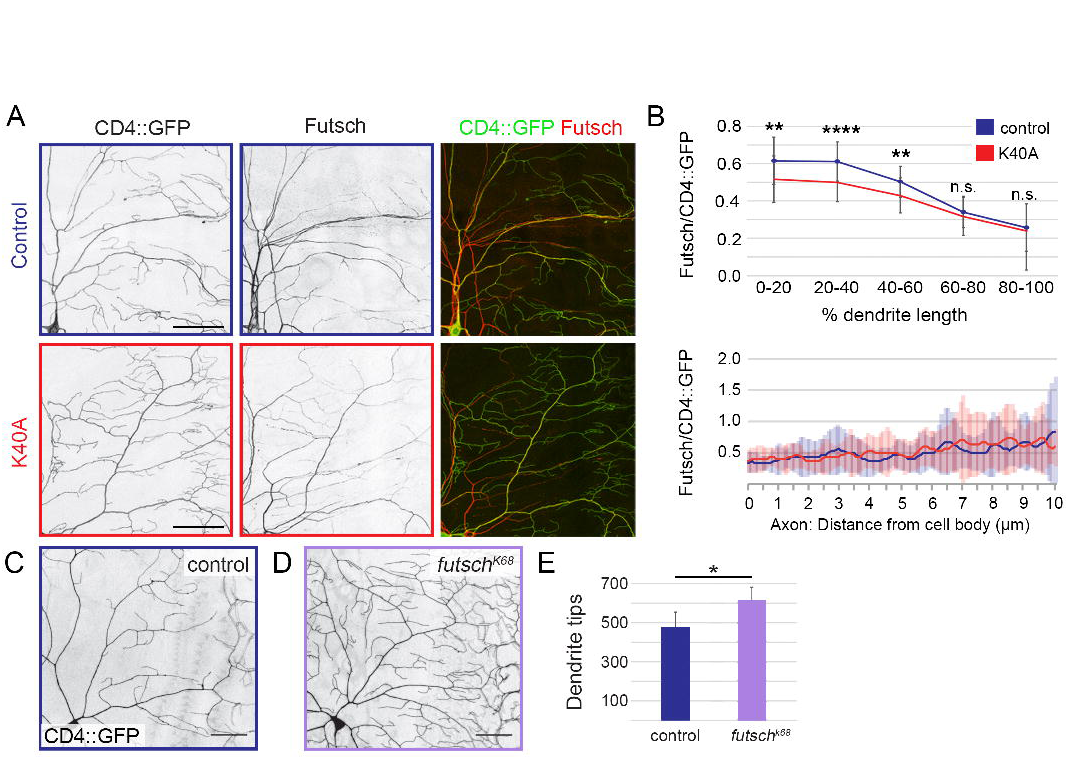
Futsch levels are decreased in αTub84B K40A mutant dendrites. (A) Representative images of a quandrant of da neuron dendrite arbors immunostained for CD4::GFP (left panel, green) and Futsch (middle panel, red). Top row: wild-type control, bottom row: *αTub84B*^*K40A*^. Scale bar: 50 µm. (B) Quantification of Futsch levels in wild-type control and *αTub84B*^*K40A*^ neurons (mean ± SD). Futsch levels measured along a dendrite (0-100% length, top) and proximal axon (bottom) were normalized to CD4::GFP levels. (C-E) In hemizygous Futsch mutant animals (D), the number of dendrite tips are increased relative to wild-type control neurons (C,E). Scale bar: 50 µm. Experiments included age-matched controls that were imaged and analyzed in parallel. *αTub84B*^*K40A*^ Futsch analysis (B, upper panel): 27 wild-type control and 45 *αTub84B*^*K40A*^ dendrites were analyzed, p=0.002 (0-20%), p=0.00004 (20-40%), p=0.001 (40-60%); 12 wild-type control and 18 *αTub84B*^*K40A*^ axons were analyzed (B, lower panel), no significant difference was detected at any 0.1 µm interval. Futsch loss-of-function dendrite tip analysis (mean ± SD, panel E): 4 wild-type control and 8 *futsch*^*K68*^ neurons were analyzed in 3^rd^ instar male larvae, p=0.01. Statistical significance was evaluated using a two-tailed Student’s t-test. ****p<0.0001, **p=0.001-0.01, n.s. = not significant.

## DISCUSSION

Acetylation of α-tubulin K40 is a highly conserved and well-studied microtubule modification. While acetylated microtubules have been shown to mediate touch sensation in invertebrates and vertebrates (Akella et al., 2010; Cueva et al., 2012; Morley et al., 2016; Shida et al., 2010; Topalidou et al., 2012; Zhang et al., 2002), a role, if any, for acetylated microtubules in neuronal morphogenesis has remained elusive. To investigate how acetylation of α-tubulin K40 might affect neuronal development we leveraged our new fly strain, which facilitates the rapid knock-in of designer *αTub84B* alleles and thus is a versatile tool for interrogating α-tubulin function in vivo. Our targeted mutagenesis of endogenous *αTub84B* points to a role for α-tubulin K40 acetylation and K40 in refining the terminal dendrite branches of developing sensory neurons. Although microtubules in young axons of cultured mammalian neurons are enriched in acetylated microtubules (Song and Brady, 2015), we found that microtubule acetylation levels are equivalent between axons and dendrites in mature sensory neurons in vivo. Mutating αTub84B K40 does not affect selective transport to axons or dendrites in these neurons, consistent with previous reports that microtubule acetylation alone is not sufficient to direct transport to either compartment (Hammond et al., 2010; Kaul et al., 2014; Witte et al., 2008). Instead, our results show that mutating αTub84B K40 alters microtubule growth, lysosome transport, and Futsch levels in dendrites but not axons. Our findings are consistent with the idea that α-tubulin K40 may be important for locally and dynamically modulating microtubule function in neurons.

Our data suggest that the increase in dendrite tips in the αTub84B K40 mutants likely reflects a change in the refinement of terminal branches that occurs during larval stages rather than an effect on dendrite outgrowth during embryogenesis. The class IV da neurons have both microtubule- and actin-rich dendrite branches (Grueber et al., 2002; Jinushi-Nakao et al., 2007), and our analyses indicate that only a subset of terminal dendrites contain acetylated microtubules (albeit dendrites with just a few microtubules might be below our level of detection). It is possible that the effect of mutating K40 on dendrite branching is modest since only a fraction of terminal dendrites contains microtubules. In contrast, mutations that disrupt the actin cytoskeleton typically produce striking changes in terminal branching (Ferreira et al., 2014; Jinushi-Nakao et al., 2007; Lee et al., 2003; Lee et al., 2015; Soba et al., 2015). One model consistent with our results and the findings of others is that microtubule acetylation fine-tunes the dynamic remodeling of microtubule-based dendrite branches. Another possibility is that mutating K40 alters the structure of α-tubulin in a way that disrupts dendrite branching.

Our data show that the αTub84B K40A mutation has modest yet significant effects on lysosome transport, microtubule growth and Futsch distribution in dendrites. While these changes might all independently contribute to an increase in dendrite branch number, it is also possible that they are mechanistically linked. We found that retrograde lysosome flux nearly doubles in *αTub84B*^*K40A*^ dendrites while anterograde lysosome transport is normal. Studies in fly and mammalian neurons indicate that MAPs, including Futsch and MAP6, can selectively disrupt anterograde or retrograde transport (Schwenk et al., 2014; Stephan et al., 2015). Notably, MAP6-regulated retrograde lysosome transport affects dendrite growth in cultured hippocampal neurons (Schwenk et al., 2014), which suggests that dendrite branching may be sculpted by the flux of lysosomes moving to and away from the cell body. It is unclear why lysosome transport, microtubule growth, and Futsch distribution are not significantly altered in the *αTub84B*^*K40A*^ axons, although compartment-specific differences in microtubule regulators and/or MAPs might make dendrites more sensitive to the αTub84B K40 mutations than axons. It is possible that several microtubule-based activities that impinge on dendrite branching are affected by mutating α-tubulin K40.

Recent studies suggest that acetylation increases the resiliency of microtubules and protects them against mechanical breakage (Portran et al., 2017; Xu et al., 2017). The da neuron dendrites, sandwiched between the larval cuticle and muscles, are potentially exposed to repeated external and internal mechanical forces. Unacetylated microtubules in *αTub84B*^*K40A*^ dendrites may be less resilient to mechanical stresses and thus may contain a higher proportion of damaged and broken microtubules than wild-type neurons. The breakage of unacetylated microtubules in worm neurons lacking αTAT1 is suppressed by paralyzing the worms (Topalidou et al., 2012). These broken microtubules are postulated to stimulate neurite branching by promoting microtubule growth from the broken microtubule ends. A similar mechanism may increase terminal branching in the *αTub84B*^*K40A*^ dendrites; however, we observe a decrease, not increase, in microtubule growth frequency. A change in microtubule flexibility and/or lattice integrity may also affect the binding of MAPs. For example, the MAP doublecortin preferentially binds curved microtubule segments, which may be prevalent in neurons with flexible microtubules (Bechstedt et al., 2014). It would be interesting to determine whether wild-type and *αTub84B*^*K40A*^ dendrites are differentially sensitive to mechanical force, and whether changes in mechanical stress would modify any of the *αTub84B*^*K40A*^ phenotypes.

An alternative interpretation of our data is that the increase in branch number is not due specifically to the loss of microtubule acetylation. For example, other modifications of α-tubulin K40 have been reported, including methylation by SetD2 (Park et al., 2016). However, it is not known whether α-tubulin K40 is methylated or otherwise modified in neurons. Lysine-to-arginine or -alanine mutations are often used interchangeably to block acetylation, although some of our results suggest that these mutations may not be entirely equivalent. This raises the possibility that intact K40 may be important to the structure of α-tubulin and/or microtubules in neurons. Consistent with this idea, work in plants has revealed that plant growth is disrupted by the expression of α-tubulin with a K40A, but not K40R, mutation (Xiong et al., 2013).

Neuronal microtubules are enriched in other α-tubulin modifications, including (de)tyrosination and polyglutamylation, whose roles in neuronal development and function are still being unraveled. We found that targeted mutagenesis of residues that are modified in the α-tubulin C-terminal tail (αTub84B Δ3 and AAAA) has no effect on animal survival. This was somewhat unexpected, given the findings, for example, that the detyrosination-tyrosination cycle affects kinesin activity (Sirajuddin et al., 2014) as well as loading dynein onto microtubules (McKenney et al., 2016; Nirschl et al., 2016), and that the loss of polyglutamylase activity alters synaptic function (Ikegami et al., 2007). However, it is important to consider that fly and mammalian microtubules may be differentially enriched in these modifications. An early report suggests that fly microtubules are only weakly detyrosinated (Warn et al., 1990). Moreover, differences in the repertoire of modifying enzymes between flies and mammals suggest that PTM dynamics may differ as well. For example, although fly microtubules are tyrosinated and detyrosinated, the lack of a known α-tubulin tyrosine ligase makes it unclear whether microtubules cycle between these two states in flies.

Our results are consistent with the possibility that PTMs may function synergistically rather than independently to regulate microtubule function (Atherton et al., 2013; Hammond et al., 2010; Kaul et al., 2014). Also, these modifications may be important to preserving microtubule-based functions in aging neurons given that changes in acetylation, detyrosination/tyrosination, and polyglutamylation have been implicated in neurodegeneration (Song and Brady, 2015). In support of this idea, we have found that adult *αTub84B*^*K40A*^ flies display an age-related deficit in righting behavior (H.R., B.V.J., J.W., unpublished data). Our current studies are not an exhaustive analysis of all known modifications of α-tubulin and microtubules. It will be of great interest to determine whether combinations of known modifications, or currently uncharacterized modifications, contribute to creating a polarized neuron. Proteomic studies have identified additional α-tubulin lysines that are acetylated (Choudhary et al., 2009; Liu et al., 2015a; Liu et al., 2015b), raising the possibility that the acetylation of other lysines might play an essential role in neuronal development.

## MATERIALS AND METHODS

### Fly strains

The *αTub84B*^*attP-KO*^ strain was created using an ends-out recombination approach (Huang et al., 2009). All *αTub84B* knock-in strains were made by using standard molecular biology methods to modify *αTub84B* in a plasmid containing an *attB* site; the plasmid with the modified *αTub84B* was then injected into *αTub84B*^*attP-KO*^ embryos expressing ΦC31 by Bestgene Inc (Chino Hills, CA). A fly strain with wild-type *αTub84B* knocked into the locus (*αTub84B*^*K'in-WT*^) rescued the lethality of the *αTub84B* knock-out. α-tubulin protein levels and dendrite branch number were equivalent between *αTub84B*^*K'in-WT*^ and wild-type flies. Thus, wild-type flies were used as controls in the experiments. The following alleles and transgenes were used in this study: *ppk-CD4::tdGFP*, *ppk-CD4::tdTomato*, *ppk-Gal4*, *UAS-Lamp1::GFP* and *Futsch*^*K68*^ (Hummel et al., 2000) were obtained from the Bloomington Stock Center (Bloomington, IN, USA); *UAS-EB1::GFP* (Rolls et al., 2007) from M. Rolls (Penn State University, University Park, PA, USA), and *UAS-katanin-60* (Mao et al., 2014) from S. Jin and Y. Zhang (Hubei University, Wuhan, Hubei and Chinese Academy of Sciences, Beijing, China, respectively). *ppk-ManII::GFP* was created by cloning *ManII::GFP* (Ye et al., 2007) downstream of the *ppk* enhancer in the *pACUH* vector (Addgene, Cambridge, MA); *ppk-ManII::GFP* was integrated at *attP VK00037* by BestGene Inc.

### Imaging and analysis

Images were acquired on a Leica SP5 laser scanning confocal microscope, equipped with two standard PMTs and a HyD GaAsP detector, using a 20X 0.7 NA oil immersion HC PL APO objective or a 40X 1.3 NA oil immersion HCX PL APO objective. Fruit fly larvae were imaged live in a drop of 50% glycerol (catalog number G153-1, Fisher Scientific, Hampton, NH) in PBS. The ratio of anti-acetylated microtubules in axons and dendrites was obtained from class IV ddaC neurons in fixed larval fillets stained with anti-acetylated tubulin (6-11-B1) and HRP. Acetylated tubulin signal in axons and dendrites was traced and captured via line scan analysis in ImageJ/FIJI (NIH) and exported to Excel (Microsoft). Signal normalized to HRP produced a similar ratio. For live neuron imaging, larvae were immobilized during imaging by pressure from a coverglass secured by two lines of vacuum grease flanking the animal. EB1::GFP and Lamp1::GFP movies were collected at rates of 1.25 frames per second (f s^−1^) and 0.5 f s^−1^ (EB1::GFP), or 1.51 f s^−1^ (Lamp1::GFP). Kymographs were generated and traced in Metamorph (Molecular Devices, Sunnyvale, CA), and the data were analyzed in Excel. Velocity (EB1::GFP, Lamp1) was calculated from the slope of the trajectory traced in Metamorph. The trajectories of lysosomes that changed speed or direction were segmented, and the segments were included in the total tallies. Lamp1 flux describes the number of lysosomes moving within an axon or dendrite segment within an approximately one-minute-long movie segment. For analyzing Futsch levels in dendrites and axons, images of neurons in fixed tissue were acquired and then dendrite segments were traced in ImageJ/FIJI using the CD4::GFP signal as a guide. Data from line scans of the CD4::GFP and Futsch signals in the proximal axon and dendrite segments were imported into Excel. We normalized anti-Futsch signal intensity by generating a ratio of Futsch to CD4::GFP. Since the dendrites included for analysis varied somewhat in length, we normalized dendrite length by dividing each dendrite into five segments that represented a percentage of the total dendrite length (e.g. the most proximal segment represented 0-20% of the total length). Dendrite tips were counted using either Imaris (automated dendrite tip counting following manual adjustments) or Metamorph (manual tip marking). Dendrite extension, retraction, and de novo growth were analyzed as previously described (Soba et al., 2015). Briefly, z-stack images of neurons expressing CD4::GFP in larvae at 48 h AEL were acquired 15 minutes apart. Maximum projections of images taken at both time points were aligned in ImageJ using the bUnwarpJ plugin and then overlaid in Metamorph. The first image (t=0) was pseudo-colored red and the second image (t=15 min) was pseudo-colored green. Overlaid images were manually scored for red and green tips using Metamorph and Photoshop (Adobe, San Jose, CA), and data were analyzed in Excel (Microsoft). All data were double-blinded before analysis and a portion of data sets were analyzed independently by two people to ensure samples were scored equivalently. Experiments were replicated at least twice.

### Immunohistochemistry

Larvae were dissected in PHEM buffer (80mM PIPES pH6.9, 25mM HEPES pH7.0, 7mM MgCl_2_, 1mM EGTA) and fixed in 4% PFA in 1X PBS with 3.2% sucrose for 45-60 minutes. The fixed fillets were then permeabilized in PBS with 0.3% Triton-X100, quenched by 50mM NH_4_Cl, and blocked in blocking buffer composed of 2.5% BSA (Sigma catalog number A9647), 0.25% FSG (Sigma catalog number G7765), 10mM glycine, 50mM NH_4_Cl, 0.05% Triton-X100. The fillets were incubated with primary antibodies overnight at 4°C in blocking buffer. Samples were washed extensively in PBS with 0.1% Triton X100 and then incubated with secondary antibodies in blocking buffer overnight at 4°C in the dark. After washing, samples were mounted on slides using elvanol with antifade (polyvinyl alcohol, Tris 8.5, glycerol and DABCO, catalog number 11247100, Fisher Scientific, Hampton, NH). Embryos were dechorionated in bleach for 1-2 min, fixed in 4% formaldehyde overlaid with heptane for 20 min, and devitellinized by rapid passage of embryos through a heptane:methanol interface. Embryos were incubated with primary antibodies diluted in PBS with 0.1% Triton X100 overnight at 4˚C and with secondary antibodies for 2.5 hours at room temperature (following each antibody incubation step, embryos were washed 3X 20 min with PBS with 0.1% Triton X100 at room temperature). Antibodies used: anti-acetylated tubulin 6-11B-1 (1:1000, or 1 µg mL^−1^, catalog number T6793, Sigma-Aldrich, St. Louis, MO), anti-polyglutamylated tubulin GT335 (1:1000, gift of Carsten Janke), anti-tyrosinated tubulin YL1/2 (1:250, or 4 µg mL^−^1, AbD Serotec MCA77G, Bio-Rad, Hercules, CA), anti-Futsch 22C10 (1:50, Developmental Studies Hybridoma Bank, Iowa City, IA), anti-HRP conjugated Alexa Fluor 647 (1:1000, or 1.4 µg mL^−1^, catalog number 123-605-021, Jackson ImmunoResearch, West Grove, PA), Dylight 550 anti-mouse (1:1000, or 0.5 µg mL^−1^, catalog number SA5-10167, ThermoFisher, Waltham, MA),.

### Immunoblotting

For western blot analysis of tubulin expression, 10 fly heads were homogenized in 30 µl of 1X SDS loading buffer. Lysate from the equivalent of one fly head (3 µl) was loaded into each lane. Proteins were transferred to PVDF membranes (catalog number 162-0177, Bio-Rad, Hercules, CA) overnight and the membranes were stained with Ponceau S (catalog number BP 103-10, Fisher Scientific, Hampton, NH) to check for efficient transfer. Membranes were blocked (5% milk, TBS Tween-20) for 1-2 hours at room temperature and incubated with primary antibody overnight. After washing, membranes were incubated with secondary antibody for 2-4 hours at room temperature. The membranes were then imaged with either chemiluminescence (SuperSignal West Pico, catalog number 34077, ThermoFisher, Waltham, MA) or fluorescent imaging using an Odyssey Imaging System (Li-Cor Biosciences, Lincoln, NE). Fluorescence intensity ratios obtained from the Odyssey were analyzed using FIJI and Excel. Antibodies used: anti-alpha-tubulin DM1A (1:1000, or 1 µg mL^−1^, catalog number T6199, Sigma-Aldrich, St. Louis, MO), anti-acetylated tubulin 6-11B-1 (1:10000, or 0.1 µg mL^−1^, catalog number T6793, Sigma-Aldrich, St. Louis, MO), anti-tyrosinated tubulin (1:1000, 1 µg mL^−1^, AbD Serotec MCA77G, Bio-Rad, Hercules, CA), anti-polyglutamylated tubulin (1:4000, catalog number T9822 Sigma-Aldrich, St. Louis, MO), and anti-actin (1:5000, Chemicon MAB 1501, EMD-Millipore, Billerica, MA).

### Statistical analysis

Multiple comparisons were performed using one-way ANOVA with post-hoc two-tailed Student’s t-tests between experimentally matched control and mutant samples. Two-tailed Student’s t-tests were used to compare two conditions. *p=0.05-0.01, **p=0.01-0.001, ***p=0.0001-0.001, ****p<0.0001, n.s.=not significant. Errors bars indicate standard deviation.

## Acknowledgements

We thank Yuh Nung Jan (University of California, San Francisco, CA) for generously supporting the initial stages of this research. We thank Drs. Yongqing Zhang and Shan Jin for katanin-60 reagents, the Developmental Studies Hybridoma Bank (created by the NICHD of the NIH and maintained at The University of Iowa, Department of Biology, Iowa City, IA 52242) for anti-Futsch 22C10 antibody, the Bloomington Drosophila Stock Center (NIH P40 OD018537) for stocks, Kevin Eliceiri and the Laboratory for Optical and Computational Instrumentation for image analysis support, and the Wickens lab for sharing reagents. We thank Jay Parrish (University of Washington), Melissa Gardner (University of Minnesota), and Wildonger lab members for discussions and comments on the manuscript.

## Author Contributions

J.W. and B.V.J. conceived of the project and wrote the manuscript. B.V.J., H.A.J.S., H.R., D.M.J.S., and J.W. performed the experiments and completed the data analysis. All authors discussed the results and provided input on manuscript.

## List of Symbols and Abbreviations

PTM: post-translational modification
da: dendritic arborization
ppk: pickpocket
h: hours
AEL: after egg laying
MAP: microtubule-associated protein

---

**Supplemental Fig. 1. Dendrite branching is similar between neurons over-expressing katanin-60 in wild-type and αTub84B K40A animals**. (A-C) Dendrite tip number is not significantly different between neurons over-expressing *katanin-60* in a wild-type versus *αTub84B*^*K40A*^ background. Scale bar: 100 µm. Experiments included age-matched controls that were imaged and analyzed in parallel. Dendrite tip analysis: 9 neurons over-expressing *katanin-60* in a wild-type background and 7 neurons over-expressing katanin-60 in a *αTub84B*^*K40A*^ mutant background were analyzed (mean ± SD). Statistical significance was evaluated using a two-tailed Student’s t-test. n.s. = not significant.

